# Response of benthic macroinvertebrates to dam removal in the restoration of the Boardman River, Michigan, USA

**DOI:** 10.1101/2020.12.22.423935

**Authors:** David C. Mahan, Joel T. Betts, Eric Nord, Fred Van Dyke, Jessica M. Outcalt

## Abstract

Dam removal is an increasingly important method of stream restoration, but most removal efforts are under-studied in their effects. In order to better understand the effects of such removals on the stream ecosystem, we examined changes in stream macroinvertebrate communities from 2011-2016 above, below, and before and after the October 2012 removal of the Brown Bridge Dam on the Boardman River in Michigan (USA), and to new channel sites created in its former reservoir (2013-2015). Using linear mixed-effect models on the percent abundance of ecologically sensitive taxa (% Ephemeroptera, Plecoptera, Trichoptera (EPT)), total density of all macroinvertebrates, and overall taxa richness, along with multivariate analyses on the community matrix, we examined differences in community composition among sites and years. EPT declined downstream of the dam immediately after dam removal, but recovered in the second year, becoming dominant within 2-4 years. Downstream sites before removal had different community composition than upstream sites and downstream sites after removal (p<0.001), while upstream and downstream sites after removal converged towards similarity. New channel (restored) %EPT, density, and taxa richness were not different from upstream sites in any year following removal, but new channel sites were the most distinct in community composition, possessing multiple indicator taxa characteristic of unique new conditions. The invasive New Zealand mud snail (*Potamopyrgus antipodarum*) was absent from all sites prior to dam removal, but appeared at low densities in upstream sites in 2013, had spread to all sites by 2015, and showed large increases at all sites by 2016. Managers employing dam removal for stream restoration should include post-removal monitoring for multiple years following removal and conduct risk analysis regarding potential effects on colonization of invasive invertebrate species.

## Introduction

Watershed fragmentation is a global ecological problem and a pervasive concern in restoration because of its detrimental effects on native aquatic biodiversity [1]. Human-created dams are one of the most prevalent causes of such fragmentation as well as altered stream flow, with more than 50% of the world’s large river systems now affected by dams and their associated impoundments [2]. Further, dams are increasingly recognized as harmful to aquatic ecosystems by acting as a strong ecological “press disturbance,” exerting constant and stressful effects on adjacent downstream and upstream reaches of the affected stream [3]. Dams alter the downstream flow of nutrients, sediment, and organic matter, and limit macroinvertebrate drift [2,3]. Dams also create barriers to movement of aquatic organisms to upstream habitats, an effect most detrimental to fish populations, particularly diadromous fish species, because dams block these species’ essential reproductive migrations and stop movement of juveniles to habitats required for optimal feeding and growth [4,5]. Additionally, dams reduce lateral connectivity in floodplain habitats and alter floodplain function by eliminating flooding [6]. As a consequence of interrupted and slower flow rates, dams change proximate upstream areas from lotic to lentic environments [4]. Cumulatively, these changes affect stream ecosystems by altering fish and macroinvertebrate habitat and trophic relationships, thus affecting community composition [7-9].

Worldwide, the United States is among the countries most affected by dams. More than 87,000 dams were in place on US waterways in 2013, with about half having been constructed between 1950 and 1980 [10]. Today many such dams are outdated, structurally unsound, not economically viable, and perceived to negatively impact stream quality [1,11] and 85% of US dams were expected to have exceeded their operational period by 2020 [12]. Such aging dams grow increasingly less functional due to accumulation of sediment behind the dam and increased danger of breaching due to structural and material deterioration [1].

There are multiple techniques appropriate to restore deficiencies created by dams and other forms of habitat-altering disturbances in river ecosystems [13,14]. The most drastic, radical, and basin-wide restoration technique is removal of the dam itself. Dam removal differs from more site-specific techniques not only in the pervasiveness of its effects on the aquatic system, but its invariable interaction with socioeconomic and political considerations that will be engaged because of the effects that dam removal may have on flood control, municipal water supply, irrigation, riparian home and property value, and power generation. Despite these multidimensional effects, dam removal has become more frequently used and pursued as a viable, often preferred strategy for stream and river restoration [10,15]. One reason that dam removal is increasingly used in river restoration is because its implementation directly addresses structural deficiencies and potential hazards of deteriorating dams. Over 1,200 dams have been removed in the United States since the 1970s, more than half of which were demolished in the last two decades [1,10,15]. However, the effects of dam removal have continued to be under-studied [1,15], with only 139 (<1%) of these dam removals accompanied by any ecological or geomorphic assessments. Such lack of scientific assessment is troubling given the increasing frequency of dam removal and its justification in terms of favorable ecological results, especially since some aspects of the less frequently assessed ecological responses have shown wide variation in magnitude and time span and may take years to detect [1,10,15].

One taxonomic group consistently used in a diverse array of aquatic habitat and stream quality assessments is benthic macroinvertebrates. Benthic macroinvertebrates are foundationally important to the food chain in aquatic systems. Because many macroinvertebrates feed on primary producers, yet are themselves consumed by organisms of higher trophic levels, they have vital ecological significance and can be influenced by both environmental pressures and biotic interactions [16]. Macroinvertebrates are sensitive to habitat quality, water pollution, and sediment changes, and many are site-specific in such sensitivity because of relatively low mobility in the stream. Therefore, changes in their abundance can serve as fine-scale indicators of stream quality, stream restoration, and recovery after disturbances [13,17-20].

Release of sediment, upstream downcutting, and change in water temperature following dam removal affect composition of macroinvertebrate communities at least in the short term [12,19,21], often resulting in an initial decline in macroinvertebrate density, especially of more environmentally sensitive taxa [22]. Following initial disturbance, dam removal has been shown to be an effective means of increasing benthic macroinvertebrate diversity and abundance of environmentally sensitive macroinvertebrates [23,24]. Just as there have been relatively few ecological assessments of dam removal, there have also been only a limited number of studies of effects of dam removal on macroinvertebrate communities, especially studies lasting more than three years [15]. As a result, there remains a continuing need for longer-term studies on the persistence of ecological responses observed in short-term studies [15], with particular value in longer-term studies of responses of benthic macroinvertebrate communities [23].

One opportunity to examine longer-term effects of dam removal arose in the Boardman River, a fifth-order groundwater-fed stream in northern Michigan, USA. As a principally groundwater-fed stream, the Boardman has an average annual discharge of 111 cfs, a 1.96 ratio of average high and low flow, and relatively stable temperature regime, making it one of Michigan’s most stable rivers from 1997-2005 [25]. There has been an ongoing effort to remove three earthen dams on the Boardman River and alter a fourth near the river’s termination into Lake Michigan via Grand Traverse Bay. The dams were created in the early 1900s to generate electricity, but are now outdated and fail to provide sufficient power to be economically sustainable [26]. The goal of these dam demolitions was not only to remove an economic burden, but to also restore natural habitat in the river and improve the recreational fishery. In addition, local Native American tribal nations supported the project as an expression of restoration and reestablishment of their own cultural heritage [27].

Dams were successfully removed in 2013, 2017, and 2018. Removal of the first and most upstream of these, the Brown Bridge Dam (BBD), was initiated in September 2012 and completed in January 2013. Restoration efforts above the dam and near the new stream channel included removal of 260,000 cubic yards of sediment, alignment with the relic stream bed, placement of more than 6,000 linear feet of woody debris for bank stabilization and in-stream habitat, downstream sediment traps, and riparian plantings [26]. Recognizing that dam removal can have a wide range of initial to longer-lasting impacts on aquatic communities [21], we used the opportunity created by the removal of the BBD to conduct a nine-year study from 2008 to 2016, monitoring responses of benthic macroinvertebrates. We used benthic macroinvertebrates as indicators of ecosystem quality to assess the river before and after the dam was removed in September 2012.

Our objective in this study was to monitor changes in benthic macroinvertebrate community composition in the Boardman River above and below the BBD, before and after dam removal, and in newly created restored channel habitat following removal. Our hypothesis was that the BBD decreased suitable habitat for native stream macroinvertebrates in the Boardman River, and that, although removal would initially disturb the system, absence of the BBD would eventually increase similarity between upstream and downstream communities of macroinvertebrates as habitat conditions changed after the removal. The fundamental research questions we addressed were: (1) what differences existed above and below the BBD prior to its removal; (2) how did removal of the BBD affect the composition of these communities in both short- and long-term responses, and (3) did macroinvertebrate communities above and below the BBD converge toward similarity after dam removal? Through our investigation, we determined how stream macroinvertebrate communities changed over time in response to the presence and removal of a dam.

## Materials and methods

### Study sites

From 2008 to 2016 we surveyed macroinvertebrate communities in 2 to 11 sites along the Boardman River (Fig 1, Table 1). Eight non-impounded sites were sampled prior to removal of the BBD for 1-4 years to determine reference conditions to which post-dam removal assessments could be compared. These sites continued to be monitored for four years after dam removal from 2013-2016 to assess post-removal response of macroinvertebrates (Fig 1, Table 1), although the furthest downstream site (LP) stopped being sampled in 2014 due to downcutting impacts related to the drawdown of the reservoir associated with another dam, the Keystone dam. Previously impounded sites above the BBD also were monitored for three years after dam removal from 2013-2015. Sites were selected based on presence of representative riffle habitat with consideration of sampling accessibility. Sites were similar in channel width, slope and forested riparian habitat, except for new channel sites where riparian habitat was not forested. An annual proposal describing the objectives and methods used in this study, including capture and preservation of macroinvertebrates, was reviewed and approved by the Adam’s Chapter of Trout Unlimited, the Conservation Resource Alliance, and the Grand Traverse Band of Ottawa and Chippewa Indians in each year of the study. Because there was no capture, handling, or mortality of vertebrates associated with the study, no additional approvals were required. No protected species were sampled. Public access to the Boardman River was at sampling sites adjacent to dams or roads. Access to the Boardman River at sampling locations adjoining private property was given by the respective private property owners.

**Table 1.** Description of study sites in riffle macroinvertebrate survey, 2008-2016, Boardman River, Michigan, USA. Sampling was conducted in early June of each year, and the Brown Bridge Dam (BBD) was removed between September 2012 and January 2013. Distance from the BBD location in meters along the river, negative values indicate distances upstream of the dam, and positive values indicate distances downstream of the dam. Only sampling in years 2011-2016 were included in analysis. The Brown Bridge Road (BBR) site was not included in analysis in 2011.

| Study site | Site Category | Dist. Dam* | Location | Years sampled |
| --- | --- | --- | --- | --- |
| Grasshopper Upper (GHU) | Upstream | -6100 | 44.652 N, -85.470 W | 2011-2016 |
| Grasshopper Lower (GHL) | Upstream | -5260 | 44.649 N, -85.476 W | 2011-2016 |
| Brown Bridge Upper (BBU) | New Channel | -4180 | 44.649 N, -85.487 W | 2013-2015 |
| Brown Bridge Middle (BBM) | New Channel | -3470 | 44.649 N, -85.490 W | 2013-2015 |
| Brown Bridge Lower (BBL) | New Channel | -182 | 44.647 N, -85.505 W | 2013-2015 |
| Brown Bridge Road (BBR) | Downstream | 1060 | 44.638 N, -85.517 W | 2012-2016 |
| 14 Ponds (14P) | Downstream | 2350 | 44.639 N, -85.529 W | 2012-2016 |
| Wyrwas (WYR) | Downstream | 6280 | 44.646 N, -85.554 W | 2012-2016 |
| Summers (SUM) | Downstream | 8610 | 44.648 N, -85.574 W | 2012-2016 |
| Shumsky (SHU) | Downstream | 11240 | 44.651 N, -85.591 W | 2008-2016 |
| Lone Pine (LP) | Downstream | 20460 | 44.685 N, -85.627 W | 2008-2014 |

**Fig 1.**
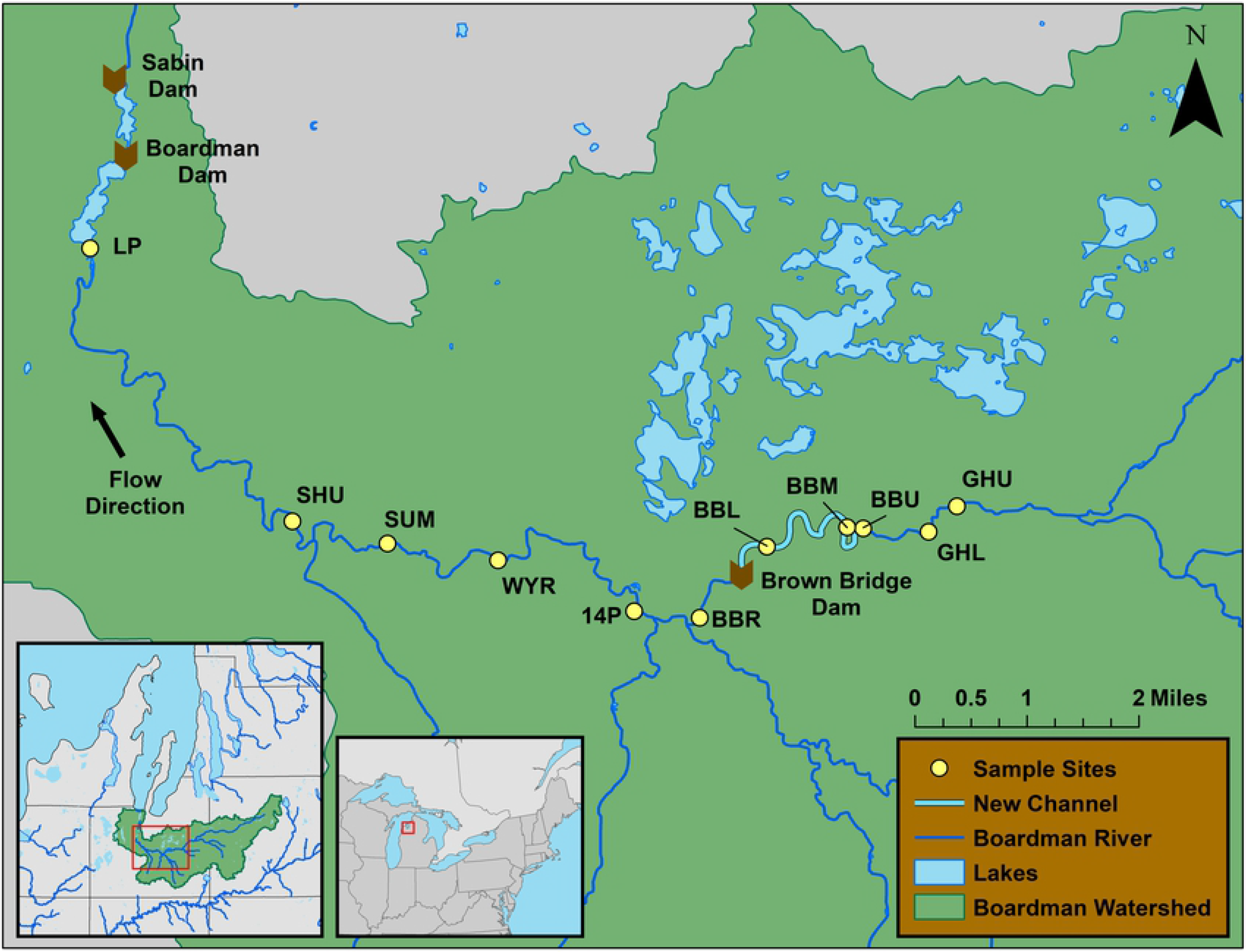
Location of study sites and Boardman River dams for riffle macroinvertebrate survey, 2008-2016, Boardman River, Michigan, USA. See Table 1 for full names and locations of sites. The Brown Bridge, Boardman, and Sabin dams were removed in 2013, 2017, and 2018, respectively. Reservoirs are pictured for later-removed dams were removed after the study period.

During the removal process there was a failure of the dewatering structure associated with the BBD on 6 October 2012. The reservoir drawdown, which had been planned to proceed gradually over 20 days, occurred in a period of six hours as a result of a site-specific weakness (“boil”) in the lining of the temporary water control structure [28]. The flow of water through the boil increased rapidly and uncontrollably, allowing water to rush downstream in an above-bank flood, creating a sudden large movement of water and sediment that inundated 66 private property holdings [28]. The dewatering event lowered the 69 ha holding pond 4.3 m during the six hour period and was subsequently estimated to have moved between 4,400 and 5,800 cubic meters of sediment downstream of the dam breach during that period [28], creating a sudden ecological pulse event intensifying dam removal impacts and possibly temporarily eradicating some macroinvertebrate species at downstream sites.

### Sampling and sorting methods

We sampled selected sites in early June each year to reduce biases associated with seasonal variation in macroinvertebrate abundance. To keep samples consistent across the length of the river, we sampled only riffle habitats using a 500 μm mesh Surber sampler with a sampling area of 0.093 m^2^ [29], with six replicates in a transect across the width of the channel, with sediment cover ranging from ∼0.25 cm to ∼10 cm and macrophyte cover of ≤5%, to reduce variation in samples [30]. For each replicate, we agitated the gravel for one minute with a pronged agitator and by hand. Organic material, substrates, and macroinvertebrates were rinsed into the bottom of the net and washed into a sample jar following sampling. The Surber sampler was then visually inspected to ensure that no organisms were lost. At each site, we recorded GPS coordinates for accuracy of location replication in subsequent years.

Following sampling, we immediately placed each replicate in a sample jar and preserved in ≥50% ethanol until sorting. We subsequently sorted and identified macroinvertebrates to family, with insects except for chironomids identified to genus if possible [31-33]. All identified macroinvertebrates were then preserved in 95% reagent ethanol.

## Data analysis

Abundances of each taxonomic group from all replicates in a given site were summed to provide a cumulative sample with an area of 0.558 m^2^ (0.093 m^2^ × 6) for each site × year. From these samples, we determined the proportion of environmentally sensitive organisms present using %EPT [3], densities (number/m^2^) of total macroinvertebrates, and taxa richness at the family level (S1 Table). We also determined the density of total macroinvertebrates in each functional feeding group [34], including collector/filterer, collector/gatherer, parasite, predator, scraper, and shredder feeding groups (S1 Table). Due to disproportionately high abundances of hydrobiid snails in 2016, this taxon was removed from all of the above calculations except taxa richness. To identify trends in macroinvertebrate communities over time, we noted the three most abundant families at each site. In cases where there were equal abundances for any of these families, the next most abundant family was also included.

To create comparisons that would address our research questions, site × year designations classified in one of four site categories in relationship to the removed dam: sites upstream from the BBD (upstream), sites downstream from the BBD before removal (downstream-before), sites downstream from the BBD after removal (downstream-after), and sites within the impoundment created by the BBD (new channel sites). We further categorized sites according to our assessment of the impact of dam removal on such sites—strongly impacted (downstream-after) and weakly impacted (downstream-before and upstream) for comparison in linear models.

We tested for evidence of dam removal impact on the summary variables described above (%EPT, macroinvertebrate density, and taxa richness) and on macroinvertebrate functional feeding groups (FFGs) using linear mixed-effect models fitted with removal impact class or site × category as a fixed effect and site and year within site as random effects. To investigate year-to-year differences, the same response variables were modeled with removal impact class × year interactions as fixed effects and site as a random effect. Functional feeding groups were not modelled with removal impact class × year interactions. Parasites were not included in FFG analysis because they were represented by only one taxa (Gordiidae). All models were fitted using the function *lme* from package *nlme* [35] and post hoc comparisons were evaluated for significant linear models using the function *glht* from package *multcomp* [36] in R version 3.3 [37]. Linear regression was used to test for competitive exclusion between hydrobiid snails and other scraper taxa.

We further explored similarity of the benthic communities across site × years using PERMANOVA to test whether the groups defined by the effects of dam removal or spatial relationship to the dam were distinct from one another in multi-dimensional space (i.e., whether macroinvertebrate community composition differed between site categories). PERMANOVA is a multivariate analog of the more commonly used analysis of variance (ANOVA) where the total dispersion of an n-dimensional dataset is partitioned into dispersion of group centroids from the overall centroid and residual dispersion (dispersion around group centroids). Permutation analysis is used to calculate *p*-values based on an F-ratio of the dispersions to test the significance of cluster differences [38]. Non-metric Multi-Dimensional Scaling (NMDS) was used to visually represent these differences. We fitted the PERMANOVA and NMDS to fourth-root transformed community data using Bray-Curtis distance [38] and 10^6^ permutations for the p-value, using the package *vegan* [39]). Hydrobiid snails were not included in this analysis because the dominance of this group diluted other potentially important findings.

We followed NMDS and PERMANOVA with SIMPER and Indicator Analysis to determine which taxa were most significantly associated with the differences in community composition between each impact category and to organize taxa by feeding groups. We used the 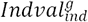 procedure [40] on fourth-root transformed invertebrate densities. Analysis was carried out with the *multipatt* function with *IndVal*.*g* in package *indicspecies* R version 1.7.6 [41]. For NMDS, PERMANOVA, SIMPER, and Indicator Analysis, rare taxa with ≤5 individuals in the in the entire data set were excluded [42].

## Results

### Distribution of macroinvertebrate families

Sampled sites varied in abundance of macroinvertebrate families by site and year. Among 58 samples from 11 sites over a nine-year period, the most commonly present taxa were Chironomidae (*n*=57 samples), Ephemerellidae (*n*=56), Baetidae (n=54), Brachycentridae (n=54), Elmidae (*n*=54), Athericidae (n=50), and Simuliidae (n=47). The most dominant taxa were Ephemerellidae (*n=*37), Chironomidae (*n*=32), Elmidae (*n*=31), and Hydrobiidae (*n*=23) (where *n =* # of times occurring in three most abundant taxa at a site; S2 Table). With the exception of 2013, we observed more dominance of environmentally sensitive taxa (EPT families) at all sites × years following dam removal, along with a large increase of New Zealand mud snail (*Potamopyrgus antipodarum*) abundance in 2016, and a decrease in dominance of families Chironomidae and Simuliidae over time following dam removal (S2 Table). Seven months after dam removal, in 2013, chironomids, simuliids, and/or aquatic worms (Oligochaeta) were present in relatively high numbers at almost every downstream and new channel site. However, we did not observe chironomids or simuliids in the three most abundant families at any site in our final year of sampling in 2016.

### Differences by site category, dam removal impact class, and year

When considering sites grouped as site categories (upstream, downstream-before, downstream-after, and new channel) and not considering the effect of year, there were no differences in invertebrate density (F_[3/43]_=1.67, *p=*0.19), taxon richness (F_[3/43]_=0.65, *p=*0.58), or %EPT (F_[3/43]_=1.33, *p*=0.28) (Table 2). Likewise, when considering sites grouped as strongly impacted (downstream-after) and weakly impacted (downstream-before and upstream) and not considering the effect of year, there were no differences for any summary variable (F_[1/39]_=1.66, 0.101, & 0.705, respectively; *p*>0.05; Table 2). In contrast, year effects showed significant differences in dam removal impact class, but only for %EPT. Seven months after dam removal in 2013, %EPT was depressed at downstream sites (strongly impacted sites) compared to both strongly impacted and weakly impacted sites in other years (F_[9/25]_=3.60, *p*=0.005; Table 2, Fig 2). In contrast, %EPT was not different between downstream (strongly impacted) and upstream (weakly impacted) sites in 2013 (Z=1.54, *p*=0.87; Table 2B, Fig 2), likely because %EPT at one of two upstream sites was also depressed (potentially affected by the drawdown in 2013). %EPT increased in 2014-2016, and was not different between strongly impacted (downstream-after sites) and weakly impacted sites (upstream sites and downstream-before sites) for any year-to-year comparison with these years except 2013 (*p*>0.25; Table 2, Fig 2). Macroinvertebrate density increased steadily at downstream sites after a low in 2013, although these trends were not significant (*p*>0.25 for all year to year comparisons; Fig 2). There was an upward trend in taxa richness from 2013 to 2016, although differences also were not significant (F_[9/25]_=1.79, *p*=0.12; Table 2A, Fig 2). New channel sites had consistently high values of macroinvertebrate density, taxon richness, and %EPT in all years including in 2013, seven months after dam removal, and were not different than upstream sites in any summary variable (Table 2Aa, Fig 2).

**Table 2.** A) ANOVA tables of linear models of summary variables Invertebrate density, Taxon richness, and EPT percent as predicted by different site categorizations. B) Post hoc tests for differences in EPT percent between strongly impacted (S) and weakly impacted (W) sites by year. All models exclude Hydrobiid snails. Significant p-values in bold. Post hoc comparisons with p>0.25 not shown.

| A: Summaries of linear models |  |  |  | B: Post-hoc tests |  |  |  |
| --- | --- | --- | --- | --- | --- | --- | --- |
| Comparison/Variable | df | F-value | *p* | Test | Estimate | Z-value | *p* |
| Strongly Impacted vs Weakly Impacted Site × Years |  |  |  | EPT percent |  |  |  |
| Invertebrate density | 1/39 | 1.66 | 0.205 | S2014 - S2013 | 0.334 | 4.398 | < **0.001** |
| Taxon richness | 1/39 | 0.101 | 0.753 | S2015 - S2013 | 0.303 | 3.797 | **0.005** |
| EPT percent | 1/39 | 0.705 | 0.406 | S2016 - S2013 | 0.304 | 3.81 | **0.005** |
| Site × Years Grouped by Site Category |  |  |  | W2011 - S2013 | 0.241 | 2.809 | 0.125 |
| Invertebrate density | 3/43 | 1.67 | 0.188 | W2012 - S2013 | 0.269 | 3.78 | **0.006** |
| Taxon richness | 3/43 | 0.659 | 0.582 | W2014 - S2013 | 0.456 | 4.16 | **0.001** |
| EPT percent | 3/43 | 1.33 | 0.277 | W2015 - S2013 | 0.345 | 3.146 | **0.049** |
| Strongly Impacted vs Weakly Impacted by Year (Removal Impact Class) |  |  |  | W2016 - S2013 | 0.283 | 2.575 | 0.217 |
| Invertebrate density | 9/25 | 0.737 | 0.672 |  |  |  |  |
| Taxon richness | 9/25 | 1.79 | 0.121 |  |  |  |  |
| EPT percent | 9/25 | 3.60 | **0.005** |  |  |  |  |

**Fig 2.**
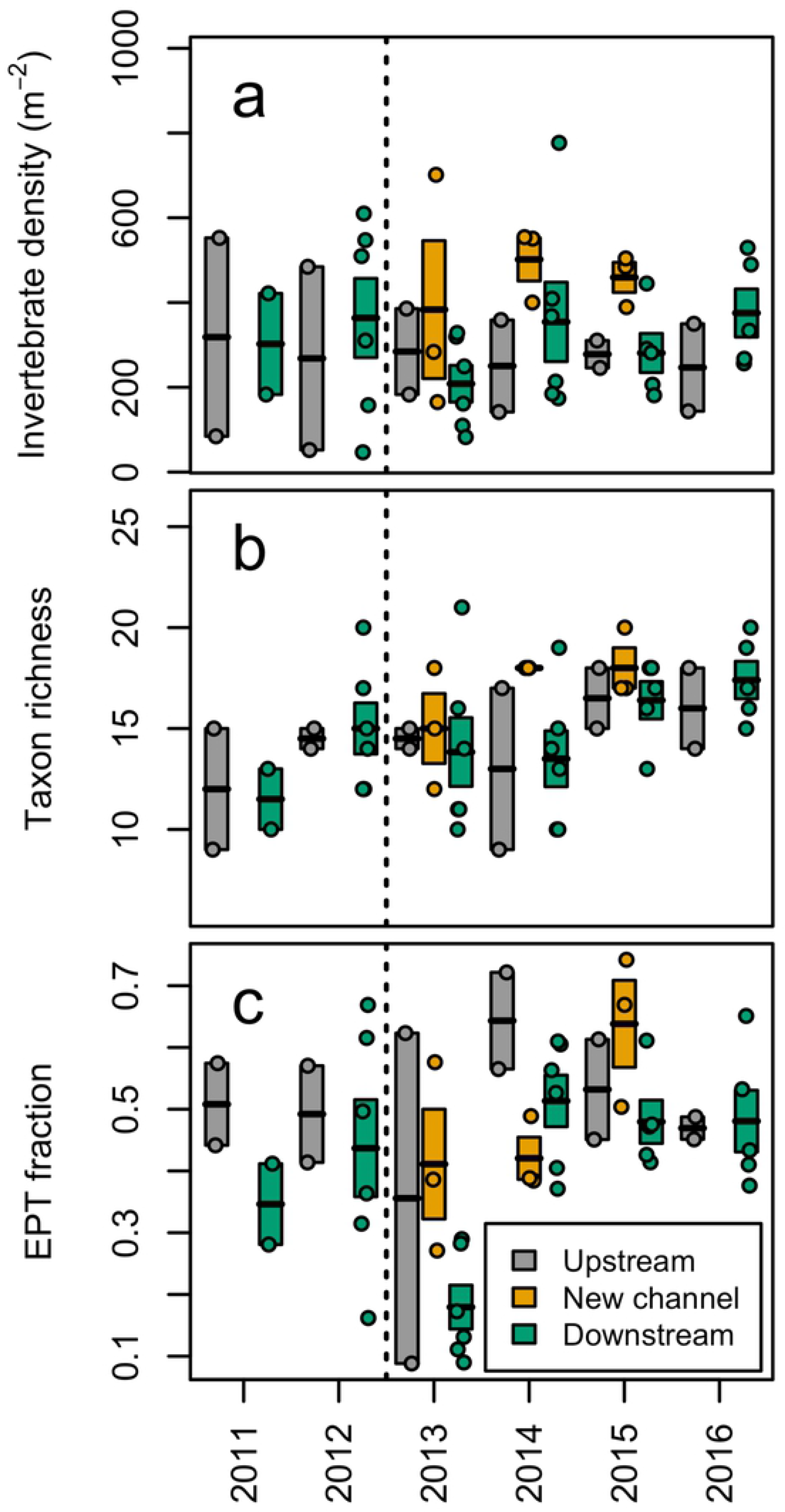
Benthic macroinvertebrate data, summed from six Surber samples (0.093 m^2^) at each of 10 sites, and averaged at upstream, downstream, and formerly impounded (new channel) sites of the Brown Bridge Dam (removed in between 2012-2013) along the Boardman River, Michigan, USA. Variables measured were: invertebrates per square meter (A), taxon richness (family level, sometimes higher classification for non-insects) (B) and fraction of insects in orders Ephemeroptera, Plecoptera, and Trichoptera (%EPT) (C). Rectangles show mean +/- SE with mean dividing each rectangle. Points indicate measured values. Dashed line indicates time of dam removal. Due to large numbers of New Zealand mud snails (*Potamopyrgus antipodarum*) at all sites in later years, this taxon was excluded from summary variable calculations.

### Community ordination

PERMANOVA analysis detected differences in community composition between site categories (F_[47]_=3.48, *p*<0.001). Pairwise PERMANOVA showed that downstream sites before dam removal had a different community of macroinvertebrates than either upstream sites (F_[1]_=4.09, *p*=0.00003) or downstream sites after dam removal (F_[1]_=4.09, *p*=0.00024).

Differences in upstream and downstream-after site categories were significant at *α* = 0.1 (F_[1]_=2.20, *p*=0.052). These differences are apparent when visualized using NMDS ordination plots with a 3-axis solution (Stress: 0.17; Linear Fit R^2^=0.83; Fig 3). New channel sites clustered together distinctly apart from other site categories, suggesting a unique community composition. PERMANOVA could not be used to compare new channel to other site categories because they were less dispersed (equal dispersion is an assumption of PERMANOVA) [38].

**Fig 3.**
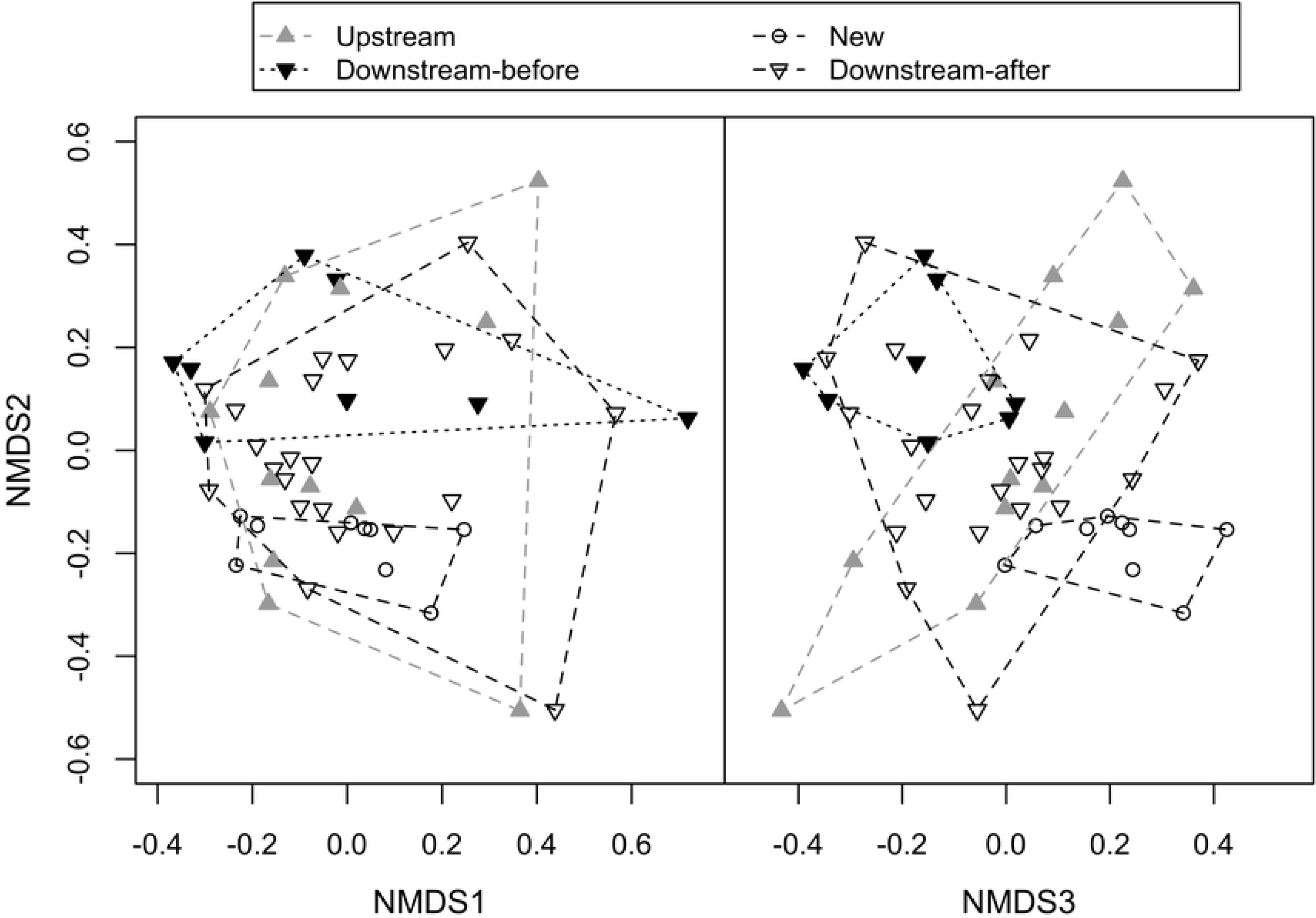
Non-metric multi-dimensional scaling (NMDS) ordination of stream macroinvertebrate community composition in the Boardman River, Michigan, (USA). 3-axis solution displayed (Stress: 0.17; Linear Fit R^2^=0.83). Ordination hulls with distinct coloration highlight site categories, Site × Years displayed with the last 2 digits of the year of sampling.

### Taxa-specific analyses

Densities of Collector-Gatherer macroinvertebrates were greater in newly restored channel sites than in all other site categories (p<0.05; Table 3). Densities of scrapers (without Hydrobiids) were higher in downstream sites before dam removal than after dam removal (p<0.05; Table 3).

**Table 3.** A) ANOVA tables of linear models for five functional feeding groups (FFG) according to Bouchard (2004) as predicted by different site categorizations. B) Post hoc tests for differences in site category comparisons for significant FFG groups (CG and SC). Post hoc comparisons with p>0.25 not shown. Significant p-values in bold. Upstream=U, Downstream-before (DB), Downstream-after (DA), New channel (N). CF = collector/filterer; CG = collector/gatherer; PR = predator; SC = scraper (without hydrobiids); SH = shredder.

| A: Summaries of linear models |  |  |  | B: Post-hoc tests |  |  |  |
| --- | --- | --- | --- | --- | --- | --- | --- |
| Comparison/Variable | df | F-value | *p* | Test | Estimate | Z-value | *p* |
| Site × Years Grouped by Site Category | | | | CG | | | |
| CG | 3/43 | 3.548 | **0.022** | N - U | 92.444 | 2.610 | **0.043** |
| CF | 3/43 | 1.970 | 0.133 | N - DB | 85.501 | 2.601 | **0.045** |
| SC | 3/43 | 3.844 | **0.016** | N - DA | 95.148 | 3.113 | **0.010** |
| SH | 3/43 | 2.590 | 0.065 | SC |  |  |  |
| PR | 3/43 | 0.0938 | 0.963 | DB - U | 48.981 | 1.978 | 0.187 |
|  |  |  |  | DB -DA | 41.484 | 3.053 | **0.011** |
|  |  |  |  | N - DB | -56.203 | -2.387 | 0.075 |

SIMPER analysis following PERMANOVA (not including new channel sites) showed that two taxa, Helicopsychidae and Elmidae, contributed most to differences in community composition between downstream-before sites and both upstream sites and downstream-after sites. Indicator analysis by site category showed that Helicopsychidae was the most significant indicator for downstream-before sites (IndVal = 0.845, *p*=0.0002), followed by Philopotamidae (IndVal=0.588, *p*=0.0084), and Brachycentridae (IndVal=0.552, *p*=0.047), suggesting that these taxa might have been most affected by dam removal (Table 4). An unknown freshwater clam (Bivalvia) was an indicator of upstream sites (IndVal=0.648, *p*=0.0024) (Table 4). New channel sites had a variety of significant indicator taxa including Chironomidae and Gordiidae (IndVal=0.576 & 0.722, p=0.0002 (both), respectively), and Perlodidae, Lepidostomatidae, Baetidae, and Ceratopogonidae (IndVal=0.642, 0.631, 0.573, and 0.583; p=0.0034, 0.0044, 0.0014, and 0.0152, respectively) (Table 4). The high number of indicator taxa for new channel sites was consistent with the distinct clustering of these sites from other site categories (Fig 3), providing further indication of distinct macroinvertebrate community composition at these sites.

**Table 4.** Taxa that are significant (p<0.05) or marginally significant (p<0.1) indicators for each site category, with their functional feeding group (FFG) [34]. Indicator analysis was carried out on fourth-root abundance data (summed from six Surber samples) at sites along the Boardman River, Michigan (USA). Significant p-values in bold. All models here exclude Hydrobiid snails. CF = collector/filterer; CG = collector/gatherer; PA = parasite; PR = predator; SC = scraper; SH = shredder.

| Site Category | Taxa | FFG | Indicator Value | $p$ |
| --- | --- | --- | --- | --- |
| Upstream | Bivalvia | CF | 0.648 | **0.0024** |
| New Channel | Chironomidae | CG | 0.576 | **0.0002** |
| New Channel | Gordiidae | PA | 0.722 | **0.0002** |
| New Channel | Perlodidae | PR | 0.642 | **0.0034** |
| New Channel | Lepidostomatidae | SH | 0.631 | **0.0044** |
| New Channel | Baetidae | CG, SC | 0.573 | **0.0014** |
| New Channel | Ceratopogonidae | PR | 0.583 | **0.0152** |
| New Channel | Sphaeriidae | CF | 0.498 | 0.0614 |
| New Channel | Asellidae | CG | 0.436 | 0.0652 |
| New Channel | Heptageniidae | SC | 0.577 | 0.068 |
| Downstream-Before | Helicopsychidae | SC | 0.845 | **0.0002** |
| Downstream-Before | Philopotamidae | CF | 0.588 | **0.0084** |
| Downstream-Before | Brachycentridae | CF | 0.552 | **0.047** |
| Downstream-Before | Glossosomatidae | SC | 0.486 | 0.0952 |
| Downstream-After | Pleuroceridae | SC | 0.426 | 0.0808 |
| Downstream-After | Curculionidae | SH | 0.459 | 0.0888 |

### New Zealand mud snails

New Zealand mud snails were absent from all sites from 2008-2012. They first appeared at seven of 11 sites in 2013 at relatively low densities, increasing their distribution to include all sampled sites by 2015, and significantly increasing their average abundance in that year (t_[11.8]_=2.52, *p*=0.027). In 2016, mud snails increased in average density per occupied site (over 1500% on average compared to 2015), reaching densities of from 542 to over 3,000 individuals/m^2^ (t_[13.4]_=9.06, *p*=0.0000; Table 5). Our 2013 record was the first observance of the species in the watershed. We found no evidence of a negative relationship (suggesting competitive exclusion) between hydrobiid abundance and abundance of other scrapers (R^2^=0.0129, *p*>0.05).

**Table 5.** Density (individuals/m^2^) of New Zealand mud snails (*Potamopyrgus antipodarum*-Hydrobidae) from 2011-2016, Boardman River, Michigan (USA). Asterisk (*) denotes significantly higher density than in previous year. *t-*tests carried out on log-transformed data comparing each year with prior year. Significant p-values in bold. There are no *t*-test results for 2011 because 2010 data not included. There are no results for 2012 because no *Hydrobiidae* were recorded in 2012 or 2011.

| Site | Site Position | Distance to Dam (m) | 2011 | 2012 | 2013 | 2014 | 2015 | 2016 |
| --- | --- | --- | --- | --- | --- | --- | --- | --- |
| GHU | Upstream | -6100 | 0 | 0 | 45 | 16 | 122 | 757 |
| GHL | Upstream | -5260 | 0 | 0 | 330 | 109 | 34 | 1398 |
| BBU | New channel | -4180 | -- | -- | 52 | 61 | 47 | -- |
| BBM | New channel | -3470 | -- | -- | 79 | 118 | 45 | -- |
| BBL | New channel | -182 | -- | -- | 9 | 47 | 100 | -- |
| BBR | Downstream | 1060 | 0 | 0 | 11 | 11 | 108 | 834 |
| 14P | Downstream | 2350 | -- | 0 | 4 | 4 | 194 | 3281 |
| WYR | Downstream | 6280 | -- | 0 | 0 | 0 | 34 | 786 |
| SUM | Downstream | 8610 | -- | 0 | 0 | 0 | 38 | 542 |
| SHU | Downstream | 11240 | 0 | 0 | 0 | 16 | 38 | 718 |
| LP | Downstream | 20460 | 0 | 0 | 0 | 0 | -- | -- |
| Mean | -- | -- | 0 | 0 | 52.9* | 38.2 | 75.9* | 1187.9* |
| SEM | -- | -- | 0 | 0 | 32.0 | 14.1 | 17.0 | 363.1 |
| $t^*$ | -- | -- | -- | -- | 3.42 | 0.53 | 2.52 | 9.06 |
| $df^*$ | -- | -- | -- | -- | 9.0 | 17.8 | 11.8 | 13.4 |
| p-value* | -- | -- | -- | -- | **0.0077** | 0.5998 | **0.0271** | **0.0000** |

## Discussion

### Relationship to previous studies

We documented disturbance due to dams’ presence, disturbance immediately following dam removal, and recovery over time after dam removal in the newly restored channel and downstream of the removed dam. These findings address our three fundamental research questions, porviding results consistent with other studies [21,22,24]. Prior to dam removal, downstream sites possessed a different community of macroinvertebrates than upstream sites, likely due to the dam’s effects on stream habitat, as has been documented in other investigations [3,14,43]. Scraper taxa (primarily Helicophychidae and Elmidae) were more abundant downstream of the dam before removal, which implied that either these taxa were more tolerant of pre-removal conditions or that the removal subsequently affected their populations for multiple years. Following dam removal, there was an initial reduction in %EPT, concurrent with declines in taxa richness and density (Fig 2). These initial depressive effects have been documented by other investigators [3,23], and may be related to increased sediment discharge associated with the BBD removal [44,45]. The dominance of sediment tolerant taxa (chironomids, simuliids, and/or aquatic worms) at downstream sites in the year after removal also provides evidence of depressive effects [33,34].

We observed increasing abundance of sensitive taxa at downstream sites by 2015-16, and recovery of EPT %, richness, and density of macroinvertebrates downstream of the dam within two to four years to levels similar to or higher than that of the upstream sites. Increased similarity in community composition between upstream and downstream-after sites after removal demonstrated a shift towards a community composition more similar to upstream sites. This convergence suggested a rapid recovery from the initial disturbance associated with the dam removal, as well as from the long-term disturbance associated with the dam, as has been observed in similar contexts by other investigators [21,23]. Newly restored channel sites had high densities, taxa richness, and %EPT (not lower than upstream sites) within the first year following removal, suggesting establishment of high-quality stream habitat in newly restored sites soon after removal and successful restoration of the formerly impounded zone by more environmentally sensitive macroinvertebrates. Collector-gatherer taxa (primarily Chironomidae and Baetidae) were most abundant at new channel sites, indicating that these taxa could rapidly colonize newly restored habitat. However, newly restored channel sites were least similar in community composition to upstream or downstream sites, suggesting distinct ecological dynamics in the restored channel system, with ongoing and dynamic colonization of macroinvertebrate communities in this novel habitat.

### The Brown Bridge Dam removal as a unique case

Because of the failure of the dewatering structure, increased sediment discharge and flow occurred in our study river at a far greater magnitude than was expected, exacerbating the effect of the dam removal. Declines of %EPT and other metrics in 2013 were likely due to this pulse disturbance, which might have been particularly devastating for less mobile stream macroinvertebrates which would have been dislodged by the sudden flood-pulse (especially individuals in the scraper feeding group, such as Helicopsychidae) or for macroinvertebrates that respire via external gills (such as Ephemerellidae), which might have been suffocated in the sudden deposition of sediment [3].

Without minimizing the importance of the unplanned sudden reservoir drawdown in this study, it is important to recognize that, in considering all site category comparisons, sites upstream of the dam were most different from downstream sites in macroinvertebrate community composition prior to dam removal, but were more similar to downstream sites after removal.

Such a pattern suggests that removal of the barrier was a more significant driver of macroinvertebrate community change than the disturbance associated with the removal process. These changes appear to have taken effect within two years, consistent with rates of change documented in other studies [3,21], even in rivers or sections of rivers with dramatic geomorphological differences [3]. In watersheds such as the Boardman River where a dam is the sole or primary anthropogenic ecological stressor of an otherwise relatively unimpacted stream, rivers have been observed to be more likely to recover to pre-stressed biological communities and do so in a shorter time than in more disturbed systems [1]. Full restoration of species richness and densities, however, may take longer [21], and the macroinvertebrate community in the Boardman River will likely continue to shift as the stream habitat changes over the long term.

Dams limit biota through impeding migration and altering habitat [5]. Such limitations and alterations are eliminated with dam removal. We observed rapid colonization of newly restored channel and below dam habitats by environmentally sensitive macroinvertebrates, which we interpret as being facilitated by removal of impediments to downstream drift and possibly other dam associated stressors, as well as creation of new habitat in the restored channel. Dam removal, then, might be an important change in the stream system ultimately leading upstream and downstream sites toward greater species and community convergence, as was the case in this study and has been the case in other investigations [3]. Other factors contributing to rapid recovery of community assemblage in the Boardman River could also include healthy upstream source populations, successful in-stream habitat restoration efforts, and the generally stable nature of this groundwater-fed stream system which moderate changes in flow and discharge.

### Study constraints

An engineered restoration of the below-dam and newly formed above-dam river channels was concurrent with the dam removal itself, and so may have affected physical habitat at the same time that sedimentation and flooding from dam removal were affecting biota. Our study, however, did not incorporate methodologies that could have separated effects of habitat restoration in the river channel from that of ecological pulse disturbance from the dam removal, and the potential for interactions between the physical stream channel reconstruction and biotic response may have affected observed responses in ways we could not detect. This deficiency was magnified by the fact that we did not collect data on abiotic features of stream habitat itself, especially those of high relevance to invertebrates, such as water velocity and water temperature, levels of organic material, or sediment movement, quantity, and type. Dams, dam removal, and channel restoration affect these variables independently and synergistically, and changes in these variables ultimately control biotic response. Such considerations are of special concern because of the unplanned sudden reservoir drawdown event when a sudden, high volume release of water undoubtedly changed the physical complexity of downstream benthic habitat. Because we did not measure these features, we had to infer associations between changes in the biota and otherwise known habitat changes (such as the sediment surge and flood reported with the dewatering event).

Our estimation of recovery time is limited to riffle habitat because we did not sample pools. Pool habitats are natural areas of increased sediment deposition and would have been disproportionately affected by the dam failure event and its sudden discharge of formerly impounded sediment above the dam. Such a rapid and large influx of sediment could have been especially devastating to invertebrate communities in pools, likely resulting in increased clogging, the effects of which could have lasted longer than in riffle habitats, and which would be one of the most important determinants of stream macroinvertebrate community composition.

Our interpretation of these results has seasonal and spatial limitations. We sampled only in early summer, although stream invertebrate communities change across and within seasons. We only measured two upstream sites over all years, the minimum needed to capture any spatial heterogeneity in upstream condition compared to downstream condition. A balanced study design (same number of sites in each site category) would have facilitated results with increased clarity and more straight forward interpretations. We were unable to achieve this study design due to limitations on sampling time and costs.

### First occurrence of the New Zealand mud snail

In 2016, we recorded high densities (500 to >3,000 individuals/m^2^) of the New Zealand mud snail, which we first detected in 2013. Although it is unknown when they were introduced to the Boardman River, New Zealand mud snails have been reported recently in other rivers in the region [46,47] after their initial detection in the Great Lakes system in 1991 [48]. We found no published studies or unpublished agency reports that provided evidence of how mud snails entered the Boardman River. However, given the popularity of the Boardman for recreational fishing and canoeing, transmission of individuals on waders or boats from neighboring invaded watersheds seems most likely [49]. Whatever the movement vector leading to introduction, we suspect that the arrival of the mud snail was inevitable, given its presence in neighboring watersheds [46,47], and not initially facilitated by the presence of dams on the Boardman River or by the dam removal process. The mud snail’s spread to downstream reaches of the Boardman River, however, almost certainly was advanced by dam removal, especially because the New Zealand mud snail is known to be an effective passive disperser [49], one that could exploit the increased stream connectivity following dam removal or by rapid movement of a large volume of water associated with the sudden dewatering event. Our study supports this explanation as we observed increasing presence of mud snails at sites further downstream from 2013 to 2016. Mud snail populations also flourished in the restored channel in the former reservoir, consistent with the fact that mud snails have been observed to thrive in disturbed areas [49].

Studies of New Zealand mud snail invasions in the western United States indicate that these snails could threaten many species in native macroinvertebrate communities [50,51]. Increasing numbers of mud snails from 2013 to 2016 in the Boardman River may have interacted with changes in community composition associated with dam removal. Mud snail invasion certainly will continue to affect the invertebrate community, especially if the high densities seen in 2016 persist or increase. For example, direct competition for periphyton could cause declines in other native scraper species, although we found no evidence of a negative relationship between hydrobiid abundance and abundance of other scrapers. Further research is needed to determine the extent of ecological impact from this invasion, but it could worsen [48,52].

## Conclusions

Our results are consistent with a large body of literature supporting dam removal as a river management and restoration technique [11, 15]. Dam removal is one of multiple approaches that can, among other beneficial effects, create increased stream habitat heterogeneity, considered by many to be the most important factor in stream biodiversity [13]. Arguably the most drastic option for stream restoration at the watershed level, dam removal presents unique risks of potentially catastrophic ecological consequences, as well as risk of significant social and political impact. However, dam removal may create more and greater differences in before and after effects than other restoration methods which may produce little result, either in short- or longer-term comparisons [14].

Monitoring effects of dam removal lags behind application of the technique, with over 60% of streams with dam removals still not monitored or monitored for fewer than two years after removal [1,15]. Managers who employ dam removal should include a post-removal monitoring protocol and ensure that such monitoring has strong financial support and organizational commitment. Our results suggest that although substantially recovered, the macroinvertebrate community was still changing even 3.5 years after dam removal, so long-term monitoring is important to fully understand ecological effects.

Given our documentation of increases in abundance and density of the New Zealand mud snail coincident with dam removal, future monitoring of recovery from dam removal should include assessment of impacts from invasive species. The potential for dam removal to facilitate the spread of invasive species to previously unaffected streams has been considered regarding invasive fish, especially Sea Lamprey (*Petromyzon marinus*), in the Great Lakes system [53,54], but often disregarding invertebrate species. Managers should conduct a risk analysis before dam removal regarding potential effects on colonization, range increase, and population increase of potential invasive species, and include the conclusions of such analysis in decision criteria regarding whether or not to remove individual dams.

## Acknowledgments

We thank the Adam’s Chapter of Trout Unlimited, the Conservation Resource Alliance, and the Grand Traverse Band of Ottawa and Chippewa Indians for financial support, and we acknowledge that field work was on Ottawa and Chippewa ancestral lands and waterways. Au Sable Institute (Michigan) provided laboratory space, equipment, and housing for research assistants. D Ippolito and P Weimerslage assisted with macroinvertebrate identification. R Keys, A Sensenig, D Proppe, and M Freake reviewed preliminary versions of the manuscript. B Dawson, A Goetz, D Guebert, N Hadley, C Hayes, K Kilmer, M LaForge, J Louwsma, J Meier, D Petry, N Sather, A Scheeres, C Shoaff, M Tennell, and D Wrinkle assisted with specimen collection, identification, and analysis. S Herbst (Michigan Department of Natural Resources), W Keiper (Michigan Department of Environmental Quality, now Environment, Great Lakes, and Energy), and S Largent (Grand Traverse Conservation District) assisted with identification of New Zealand mud snails. B Summers and J Wyrwas provided access to sampling sites through their property.

---

**S1 Table**. Macroinvertebrate summary variables by site and year. See Table 1 for site names and locations. Percent of total of macroinvertebrates in orders Ephemeroptera, Plecoptera, and Trichoptera present (%EPT), densities (total # of invertebrates/m^2^), taxa richness at the family level, and densities in each functional feeding group (# of invertebrates/m^2^), including collector/filterer (CF), collector/gatherer (CG), parasite (PA), predator (PR), scraper (SC), and shredder (SH) feeding groups. Each variable represents six Surber samples summed to a sample area of 0.558 m2 (0.093 m2 × 6) for each site × year. Densities are rounded up to the nearest whole number. All calculations exclude Hydrobiid snails.

**S2 Table**. Total abundances of three most abundant families of benthic macroinvertebrates summed from six Surber samples (0.093 m2 per sample) at sites along the Boardman River, Grand Traverse and Kalkaska counties, Michigan (USA). In cases where there were equal numbers for any of most abundant families, the next most abundant families were included. Italicized families indicate tolerant taxa (families Simuliidae and Chironomidae, and subclass Oligochaeta), and bold families indicate sensitive taxa (Ephemeroptera, Plecoptera, and Trichoptera). Superscripts were added: A = phylum Annelida, C = order Coleoptera, D = order Diptera, E = order Ephemeroptera, G = class Gastropoda, I = order Isopoda, N = phylum Nematomorpha, R = class Arachnida, T = order Trichoptera.

